# Carbon monoxide exposure stimulates growth and activity of primary producers in diverse soil ecosystems

**DOI:** 10.1101/2025.01.21.634059

**Authors:** Yongfeng Xu, Ying Teng, Jing Liao, Pok Man Leung, Shixiang Dai, Yi Sun, Wenbo Hu, Hongzhe Wang, Yanning Li, Yuqi Huang, Zhiying Guo, Xianzhang Pan, Xiyang Dong, Yongming Luo, Chris Greening

## Abstract

Carbon monoxide (CO) is both a potent poison for many aerobic organisms and a desirable energy source for diverse microorganisms. Atmospheric emissions of this gas have increased since industrialization and their levels are highly elevated in many urban and natural environments; however, it is unresolved whether elevated levels of CO at environmentally relevant concentrations are primarily stimulatory or inhibitory to soil microbial communities. Here, we showed that CO exposure minimally affects microbial abundance, richness, and composition in diverse ecosystem soils, suggesting most microbes are tolerant of this gas. Genome-resolved metagenomic profiling showed that these soils harbored diverse bacteria capable of using CO an electron donor for aerobic respiration and carbon fixation. CO stimulated the growth of several of these putative CO-oxidizing bacteria in a dose-dependent manner, especially widespread proteobacterial and actinobacterial lineages. Additionally, we found a strong relationship between CO oxidation and carbon fixation and observed an enrichment of several carboxydotrophic MAGs capable of carbon fixation in most soil types. These findings highlight that environmentally relevant CO levels do not inhibit soil microbes, but instead foster the growth of distinct carboxydotrophic bacteria, suggesting a robust soil CO sink that could help mitigate anthropogenic CO emissions and contribute to carbon cycling in terrestrial ecosystems.

## Introduction

While carbon monoxide (CO) is a potent poison for many aerobic organisms, this gas has also played an ancient and sustained role as an energy and carbon source for microbial communities^1–4^. This duality highlights the complexity of CO’s interaction with microbial life. On one hand, it can exert toxic effects, particularly through the inhibition of critical enzymes such as heme-copper oxidases and hydrogenases, thereby disrupting cellular respiration and metabolism^5^. However, it is important to note that much of the existing research on CO toxicity has been conducted using CO-releasing molecules that often involve toxic metal compounds, which may obscure the true effects of CO on microbial physiology^6^. In contrast, CO-oxidizing bacteria are key members of soil communities that support carbon cycling and enhance primary production; many of these bacteria couple this process with the fixation of carbon dioxide (CO_2_), contributing to biomass production and the formation of other organic matter^3,7,8^. Numerous studies have suggested that CO not only plays an ancient and pivotal role in abiogenesis, particularly as a key intermediate of the Wood-Ljungdahl (WL) pathway, but also maintains a vital contemporary role in supporting growth and persistence for both aerobic and anaerobic microorganisms^1,7,9^. In addition, CO oxidation by these bacteria help control the level of greenhouse and pollutant gases^10–12^. Moreover, CO, as an important reductant, carbon, and energy source, interacts with the cycles of other elements, such as nitrogen and sulfur, within global ecosystems^7,13^. However, little is known about how the complex interplay between the inhibitory and stimulatory roles of CO on an ecosystem scale^14^, and how microbial populations respond to enhanced levels of this gas.

Recent work has shown that aerobic CO-oxidizing bacteria are widespread and active members in soil ecosystems^13,15–17^. Results from genomic and metagenomic analyses have revealed that soil bacteria from at least 13 distinct phyla encode form I MoCu CO-dehydrogenases, the enzymes that allow them to aerobically oxidize CO^3,15^. Two major groups of aerobic CO-oxidizers have been extensively studied: carboxydovores and carboxydotrophs. Carboxydovores consume CO at lower concentrations for energy conservation but they require organic carbon for growth^18,19^. They are a broad group of bacteria and archaea, spanning various genera in the ubiquitous phyla Actinobacteriota^20^, Chloroflexota^21^, Deinococcota^22^, Firmicutes^22^, carboxydovores use atmospheric CO as a supplemental energy source for survival during carbon starvation, with some also using CO to support lithoheterotrophic growth^13^. In contrast, carboxydotrophs are a small but diverse group of bacteria and archaea, primarily within Proteobacteria and Firmicutes, adapted to use CO at high concentrations^18,27^. They can grow autotrophically on CO as the sole energy and carbon source or mixotrophically with other substrates; electrons from CO oxidation are transmitted through the aerobic respiratory chain for ATP production, as well as through the Calvin-Benson-Bassham cycle (CBB) to fix CO ^3,7^. For example, classical studies have demonstrated that *Bradyrhizobium japonicum* grows efficiently on high levels of CO^18,28^. Recently, we proposed that microorganisms first evolved a high-affinity form I CO dehydrogenase to thrive on low CO concentrations, after which the encoding genes were disseminated across various bacterial and archaeal genera, leading to the evolution of certain lineages that exploit elevated CO levels for growth in specific microenvironments^13^. In most soil ecosystems where CO levels are low, carboxydovores appear to be the dominant CO oxidizers compared to carboxydotrophs, though it’s unknown if these dynamics shift when CO levels are high.

How soil microbial communities respond to elevated CO has not been systematically evaluated. Typically, bacteria in most soils encounter CO at atmospheric concentrations (∼90 ppbv), using high-affinity CO-dehydrogenases to metabolize this trace energy source for growth and survival^29,30^. Through their metabolic activities, these microbes contribute to the net loss of approximately 250 teragram (Tg) of CO from the atmosphere annually ^13,31^. This process has significant ecological and biogeochemical implications, supporting bacterial productivity and diversity, particularly in oligotrophic environments^13,15,32^. Additionally, these microbes act as the second largest sink in the global CO cycle, thereby regulating levels of a toxic air pollutant and a climate-forcing gas^7^. However, certain soil environments experience elevated CO levels across space and time due to anthropogenic activity and natural sources, such as fossil fuel combustion, biomass burning and plant symbionts^33,34^. For example, wildfires significantly contribute to global CO budgets, with nearby CO concentrations in air reaching levels as high as 3000 ppm during combustion^35^. In addition, elevated CO levels (ranging from ∼0.9 - 15 ppmv) are common in urban areas^36,37^. Moreover, assays in an agroecosystem indicated that CO can accumulate to levels of up to 700 ppmv in legume roots and nodules^34^. In turn, these emissions may provide a supplemental energy source for some rhizobia, enhancing their growth and survival, which could affect the abundance of nitrogen-fixing symbionts and their competition for hosts^38,39^. Previous microcosm-based studies have suggested that soil CO oxidation is inhibited by elevated CO concentrations, but these used percentage levels of the gas that are unlikely to ever be biologically relevant^40^. No controlled study has systematically disentangled how elevated CO at biologically relevant levels influences the composition, function and activity of microbial communities^41^. This is particularly important given soil communities have been exposed to elevated anthropogenic CO emissions since industrialization^7,42^. Although human activities account for approximately half of the net atmospheric CO production, the microbial soil sink has so far helped to keep atmospheric CO levels stable^3^. It is crucial to understand whether this sink remains robust at even higher exposures.

In this study, we investigated how CO exposure at four different doses (from 0 to 1,000 ppmv) influences soil communities from diverse soils, spanning 14 agricultural, forest, wetland, grassland, and desert sites in China (details showed in Fig. 1 and Supplementary Table 1). Through longitudinal genome-resolved metagenomic and biogeochemical profiling in microcosms, we resolved how the composition, capabilities, and activities of soil communities shift with CO exposure. We show that CO exposure at biologically relevant concentrations has minimal inhibitory effects, but generally stimulates the activity and growth of CO-oxidizing bacteria, including CO_2_-fixing carboxydotrophs, with both cosmopolitan and unique lineages stimulated in each soil.

**Figure 1.**
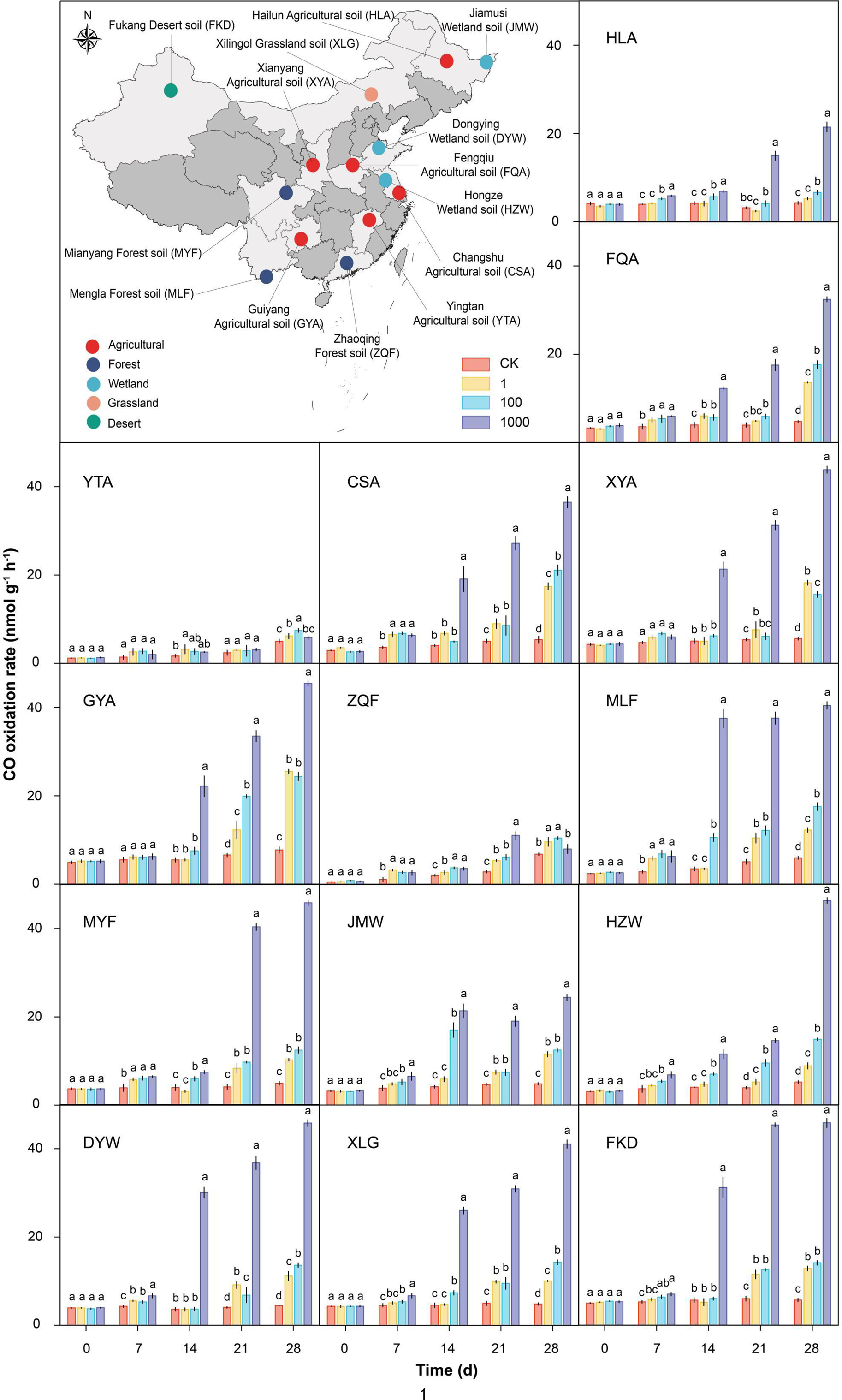
CO oxidation rate in different treatments during the soil microcosms. Map showing the distribution of sampling sites. The abbreviations refer to Hailun agricultural soil (HLA), Fengqiu agricultural soil (FQA), Yingtan agricultural soil (YTA), Changshu agricultural soil (CSA), Xianyang agricultural soil (XYA), Guiyang agricultural soil (GYA), Zhaoqing forest soil (ZQF), Mengla forest soil (MLF), Mianyang forest soil (MYF), Jiamusi wetland soil (JMW), Hongze wetland soil (HZW), Dongying wetland soil (DYW), Xilingol grassland soil (XLG) and Fukang desert soil (FKD). The designations CK, 1, 100, and 1,000 denote the different mixing ratios of CO that each microcosm was treated with (in ppmv), with CK indicating a concentration of 0 ppmv. The bars represent the means ± SD (n = 3) from three independent biological replicates. Different letters indicate significant differences between different treatments (*p* < 0.05, One-way ANOVA with Duncan’s test).

## Results and discussion

### CO exposure minimally affects soil communities but stimulates specific microbes

We first employed the 16S rRNA gene as a marker to assess how the abundance, alpha diversity, and beta diversity of bacteria and archaea in the 14 different soils (Fig. 1) were affected by CO exposure at four different concentrations (0 ppmv (control/CK), 1 ppmv, 100 ppmv, and 1000 ppmv) over 28 days. Similarly to our recent findings on H_2_ treatment on soil microbes^41^, elevated CO did not lead to significant changes of overall soil microbial community richness or abundance relative to the control, based on total 16S rRNA gene copy number or observed richness, Chao1 richness, and Shannon diversity of amplicon sequence variants (ASVs) (Fig. 2a; Supplementary Table 2). Additionally, 28 days of CO exposure minimally affected soil physicochemical properties (Supplementary Table 3). This suggests that CO exposure at biologically relevant concentrations, up to 1000 ppmv, does not cause widespread inhibition of soil communities.

**Figure 2.**
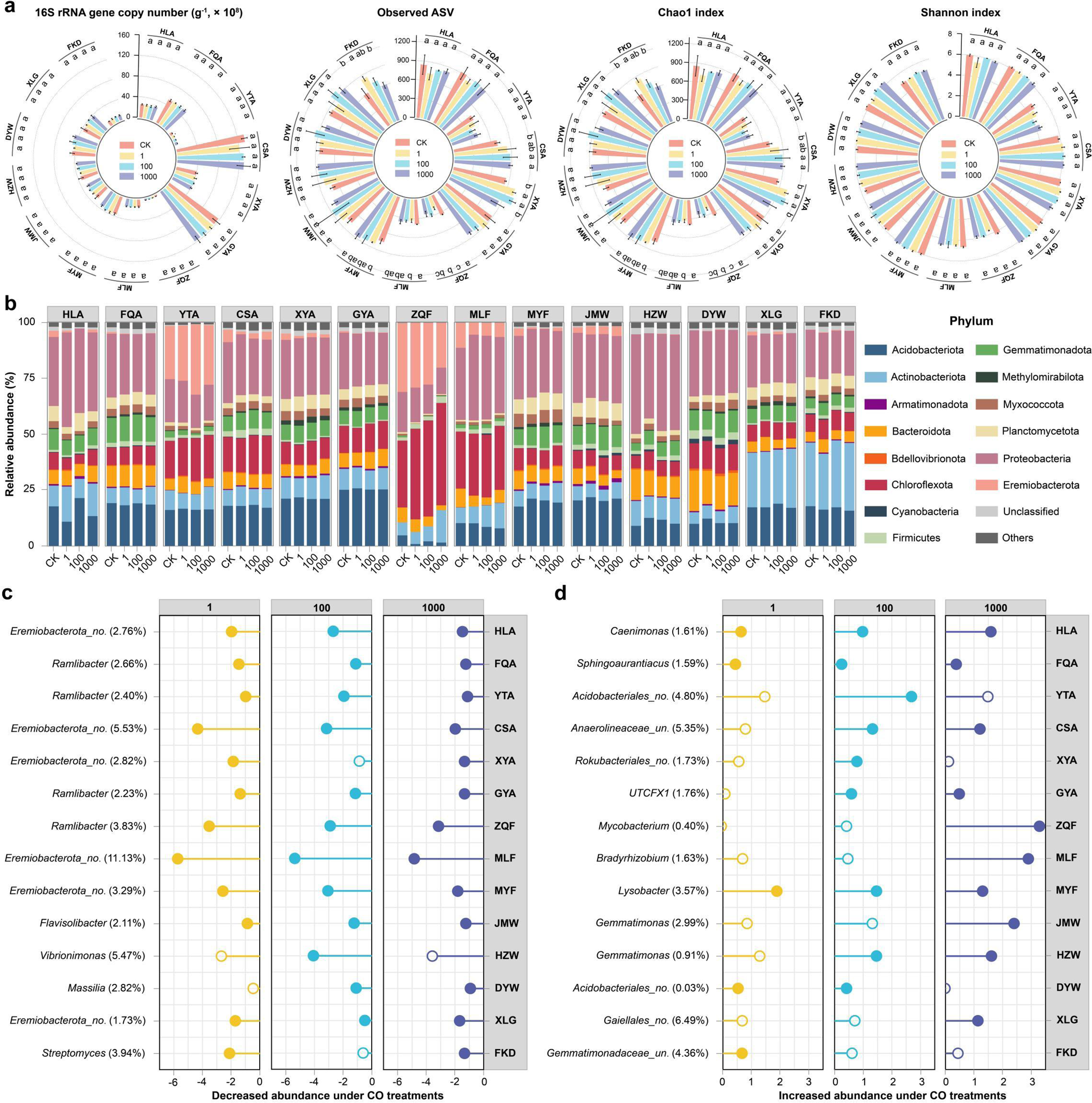
Changes of abundance, diversity, and composition of microbial communities during the soil microcosms. **a** Circular barplot of the 16S rRNA gene copy number and the alpha diversity (observed ASV, Chao1, and Shannon indexes). Different letters above the plots indicate significant differences between treatments (*p* < 0.05). **b** Stacked bar chart showing community composition at the phylum level, derived from the analysis of 16S rRNA gene amplicon sequencing and metagenomics. Phyla with abundance less than 1% in the sample were collectively categorized as “Other phyla”. **c**, **d** The most decreased and increased taxa under CO treatments in soils compared to control soils based on 16S rRNA gene amplicon sequencing analysis. The percent values in parentheses refer to the relative abundance of the phylotype in the control soil. Solid (filled) circles indicate a significant increase or decrease in relative abundance in the treatment compared to the control microcosms, with a *p*-value < 0.05; Empty circles represent *p*-values > 0.05 (One-way ANOVA with Duncan’s test).

However, CO treatment at 28 days mildly influenced bacterial community composition, with distance-based redundancy analysis (db-RDA) of beta diversity confirming CO concentration as the most significant and best predictor of changes in bacterial community composition between the microcosms (*p* < 0.05 in most ecosystem soils; Supplementary Fig. 1; Supplementary Table 4). The dominant phyla in all soils were Proteobacteria, Acidobacteriota, Chloroflexota, and Actinobacteriota, consistent with global surveys^43,44^. Significant changes in microbial community composition were observed at both phylum and genus levels in response to CO treatment. In most soils, the relative abundance of Bacteroidota and Eremiobacterota significantly decreased in the treatment vs control microcosms after 28 days, whereas Actinobacteriota were significantly promoted (Fig. 2b). However, the enriched taxa strikingly differed between the distinct ecosystem soils, with stimulated genera including the Proteobacteria *Caenimonas*, *Sphingoaurantiacus*, and *Bradyrhizobium*, as well as UTCFX1 (Chloroflexota), *Gemmatimonas* (Gemmatimonadetes) (*p* < 0.05), and various uncultured lineages (Fig. 2c; Supplementary Tables 5, 6). We hypothesized that these lineages may have been stimulated because they consumed CO, as we tested through measuring CO oxidation rates and through genome-resolved metagenomic analysis, especially given *Bradyrhizobium* is a well-known carboxydotroph^7,18,28^.

We reconstructed 630 species-level metagenome-assembled genomes (MAGs; see Supplementary Tables 7, 8) from soil metagenomes, encompassing members from 4 archaeal and 29 bacterial phyla (Supplementary Table 7). Consistent with the 16S rRNA gene amplicon profiling (Fig. 2c), most MAGs were affiliated with the phyla Proteobacteria (122), Acidobacteriota (95), Bacteroidota (65), Chloroflexota (45), Actinobacteriota (39), and Gemmatimonadota (38). Notably, we identified a considerable number of Patescibacteria MAGs (83), corroborating recent studies that indicate the prevalence of these symbionts in oxygenated soils^32,41,45,46^. In accordance with amplicon-based analyses (Fig. 2; Supplementary Table 6), certain MAGs exhibited substantial increases in abundance following elevated CO treatments, potentially due to carboxydotrophic growth. MAGs from five cultured genera (including *Microbacterium*, *Sediminibacterium, Bradyrhizobium*, and Nitrososphaera) and five candidate genera (TA-21, DTNP01, UBA10103, CADCVQ01 and a Pseudonocardiaceae genus) grew in a dose-dependent manner and sometimes became the most abundant genus in the elevated CO treatments (Fig. 3a; Supplementary Table 9).

**Figure 3.**
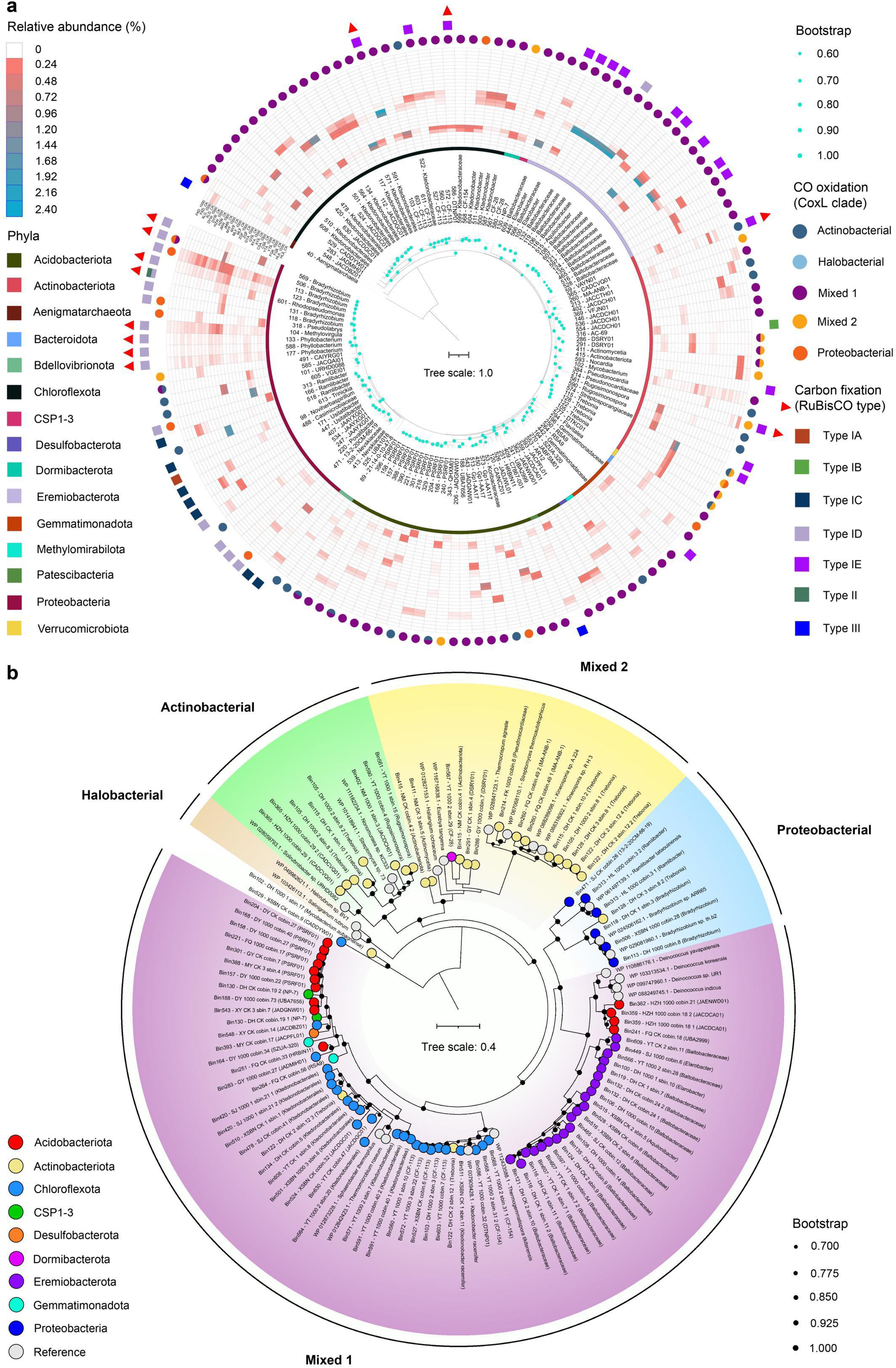
Diversity of the taxa and enzymes responsible for CO oxidation and carbon fixation. **a** Phylogenetic tree of 147 assembled soil microbial MAGs related to CO oxidation and carbon fixation. The average abundance of each MAG in the corresponding CO treatment in representative terrestrial ecosystem soils are shown in the outer circle heatmap. Taxonomy classification at the phylum level is shown in the inner circle across the 147 MAGs spanning 15 phyla. The circles indicate MAGs that encode CO-dehydrogenases and the squares indicate MAGs that encode a RuBisCO. The outermost red triangles represent the MAG that became the most abundant or were enriched in at least four ecosystem soils under elevated CO conditions. **b** Maximum-likelihood phylogenetic tree illustrating the key enzymes involved in CO oxidation. The tree displays the amino acid sequences of the large subunits of CO-dehydrogenase. The tree was rooted at mid-point while node support (1000 ultrafast bootstrap replicates) was shown in black dots. The colored leaf nodes denote phylum-level taxonomy of MAGs sequences.

### CO exposure strongly stimulates CO oxidation over time in most soil ecosystems

In parallel, we measured how the different CO exposures affected the rates of CO oxidation over time for each of the 14 sampled soils (Fig. 1). All samples consumed CO, with rates (based on averaged first-order rate constants at 1000 ppmv CO) at day 1 ranging from 0.50 nmol g^-1^ h^-1^ in a forest soil to 5.01 nmol g^-1^ h^-1^ in a desert soil (Fig. 1; Supplementary Table 10). After 28 days of incubation, CO oxidation rates were weakly affected for the control soils (CK: 4.30-7.74 nmol g^-1^ h^-1^), but strongly stimulated in the CO-exposed soils (1 ppmv: 5.26-25.51 nmol g^-1^ h^-1^, 100 ppmv: 6.65-24.39 nmol g^-1^ h^-1^, 1000 ppmv: 5.84-46.34 nmol g^-1^ h^-1^). Strong stimulation was observed for 12 soils, whereas a much weaker effect was observed for one agricultural and one forest soil (Fig. 1). Compared with the CK soil, CO oxidation rates in the 12 strongly responsive sites increased by an average of 1.2- to 3.3-fold, 1.5- to 3.9-fold and 5.0- to 10.3-fold at 1, 100 and 1,000 ppmv CO, respectively (*p* < 0.05).

Although soil CO consumption is well-documented, these findings to our knowledge the first examination of how exposure to biologically relevant CO concentrations influences CO oxidation, with most previous studies limited to very low (near atmospheric) or unrealistically high (over a million-fold higher than ambient levels) concentrations^40,47^. We also reveal that oligotrophic soils are more stimulated by CO exposure than previously thought^48^, with a 10-fold increases in CO oxidation in two wetland soils (Fig. 1). It is important to highlight that soil CO oxidation activity from nearly all sites showed a significant increase (1.2 to 3.3-fold) following treatment with 1 ppmv of CO for 28 days (*p* < 0.05; Fig. 1). This suggests that soil CO oxidizers are sensitive to even minor fluctuations in CO concentrations, at as low as 1 ppmv, highlighting the remarkable resilience of soils as effective CO sinks. It should also be noted that, even for the control soils, there was a mild and variable stimulation in CO oxidation rate after 28 days; this likely reflects that carboxydovores upregulate CO-dehydrogenase in response to nutrient starvation in order to scavenge atmospheric CO^13,21^.

### Enriched taxa encode the enzymes required for aerobic carboxydotrophic growth

We next used the MAGs to determine whether the most enriched microbes are capable of CO oxidation. Curated metabolic marker gene databases were used to annotate metagenomic data, with emphasis on the taxonomy of key genes involved in CO metabolism and carbon fixation pathways (Supplementary Figs. 3, 4; Supplementary Tables 11-14). In alignment with our recent findings from Chinese and Australian soils^15,17,41^, most community members were predicted to exhibit metabolic versatility in electron donor, electron acceptor, and carbon source usage, with the marker genes for CO oxidation (*coxL*; CO-dehydrogenase large subunit) and carbon fixation primarily through the CBB cycle (*rbcL*; RuBisCO large subunit) abundant in soils and widespread in MAGs (Supplementary Figs. 3, 4; Supplementary Table 11). As anticipated, the relative abundance of CO-dehydrogenase (specifically, Actinobacterial and Mixed 1 clades) and RuBisCO (including type ID and IE) for most soils in the elevated CO microcosms increased by an average of 5.72% (range 0.66% to 14.7%) and 1.77% (range: 0.03% to 6.7%) respectively, indicating carboxydotrophic growth. We also observed small yet significant changes in the abundance of specific genes associated with aerobic respiration, nitrogen fixation, and the oxidation of hydrogen, sulfide, thiosulfate, ammonia, and arsenite. These results underscore the significant metabolic adaptations occurring in response to elevated CO levels.

To further elucidate the genetic determinants of carboxydotrophic growth, we constructed phylogenetic trees to visualize the evolutionary relationships of CoxL (Fig. 3b) and RuBisCO (Supplementary Fig. 2) protein sequences derived from our MAGs. Aerobic CO-dehydrogenases were encoded by 122 MAGs from thirteen soil phyla, reinforcing the hypothesis of a diverse and substantial CO sink within soil ecosystems^3,7,13^. Among these, 10 Eremiobacterota, five Proteobacteria, three Actinobacteriota, and three Chloroflexota MAGs co-encoded CO-dehydrogenase alongside RuBisCO and terminal oxidases (Fig. 3a; Supplementary Tables 12-14). Nine of these MAGs increased in abundance in the CO-supplemented soils (average increase of 0.43%; range: 0.03% to 2.26%) (Figs. 3a, 4; Supplementary Table 9), including several of the aforementioned most enriched genera in CO-exposed soils. Thus, the most strongly enriched MAGs in the high CO microcosms were among those capable of carboxydotrophic growth, indicating sub-micromolar level of CO is sufficient to promote growth.

**Figure 4.**
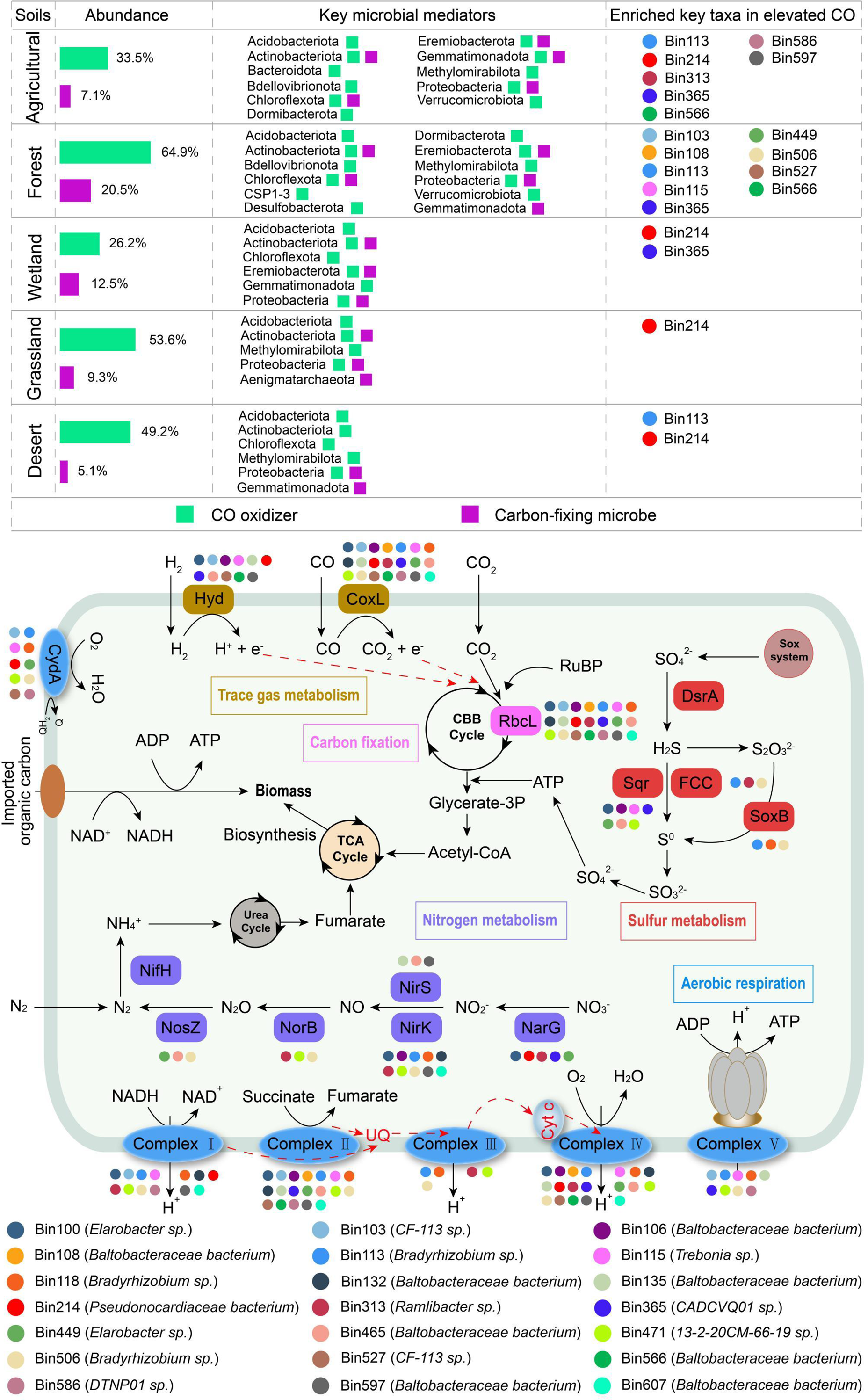
Schematic illustration of the role of distinct carboxydotrophic bacteria in different terrestrial ecosystem soils. The phylum-level affiliations and relative abundances of soil microbes that consume carbon monoxide (CO) and fix carbon dioxide (CO_2_) in the representative ecosystems are shown as bar plots. The enriched key taxa in elevated CO were showed in circles. Key taxa indicated that these MAGs encoded both CO dehydrogenases and RuBisCO. Metabolic pathways in twenty-one MAGs (with >50% completeness and <10% contamination) predicted to mediate carboxydotrophic growth.

The stimulated carboxydotrophs varied in their distributions across the sampled soils. MAGs from *Bradyrhizobium* (Bin113), a Pseudonocardiaceae genus (Bin214), CADVQ01 (Bin365), and a Baltobacteraceae genus (Bin566) were enriched in distinct biomes (Fig. 4; Supplementary Table 9). For example, a *Bradyrhizobium* MAG occurred in all soils and at least 1.1-fold stimulated in nine of them. This indicates that cosmopolitan bacteria can benefit from CO exposure to mediate carboxydotrophic growth by using electrons from CO for aerobic respiration and carbon fixation. Such metagenomic inferences are also supported by previous reports of carboxydotrophic growth in various *Bradyrhizobium* species^18,28,49^, a soil *Pseudonocardia* (Pseudonocardiaceae) isolate^50^, and the Chloroflexota phylum^51^. However, as detailed in Figure 3a and S4, some carboxydovores and carboxydotrophs showed narrower habitat ranges and were stimulated only in a specific soil. Together, these trends likely explain the common stimulation but variable rates of CO oxidation by CO exposure. Not all the enriched MAGs encoded genes for carboxydotrophy, however. For example, *Nitrososphaera* are not known to be capable of CO oxidation and instead may have been enriched due to indirect effects, whereas Patescibacteria are likely enriched as symbionts or predators of CO-enriched Actinobacteriota^15,52,53^.

### Carboxydotrophic growth of key taxa enhances carbon fixation in specific soils

Altogether, these findings suggest that certain carboxydotrophic bacteria grow in response to CO exposure by using CO as an electron donor to support carbon fixation. We therefore tested through whether CO exposure increases soil carbon fixation rates and thus primary production through microcosm-based ^13^C-CO_2_ fixation assays. Consistent with previous observations^17^, chemosynthetic carbon fixation was observed in all ecosystem soil microcosms, with average carbon fixation rates ranging from 2.3 mg kg^-1^ d^-1^ in the forest soil microcosms to 4.7 mg kg^-1^ d^-1^ in the grassland soil microcosms (Fig. 5a; Supplementary Table 15). Carbon fixation rates were significantly higher (*p* < 0.05) in eight soils from diverse biomes that received CO supplementation compared to those without CO addition, in line with the enhanced CO oxidation rates and carboxydotrophic taxa (Figs. 1, 3, 4). Conversely, we just observed a slight increase of the carbon fixation rate in three soils (HLA, FQA, HZW by 1.1-, 1.1-, and 1.2-fold) after elevated CO treatments. Although CO oxidation activity increases significantly in these soils, its carbon fixation rate does not show a significant increase. This may be attributed to the fact that some abundant carboxydovores utilize CO but do not support carbon fixation^13,21^. For example, some abundant (> 0.1%) MAGs (e.g. Bin329 in HLA, Bin218 and Bin240 in FQA, all belonging to the PSRF01 genus) harboring CO-dehydrogenases but lacking a complete CBB cycle exhibited no change in their relative abundance following treatment with high CO concentrations (Fig. 3; Supplementary Table 13). There was also no enhancement in carbon fixation in the two soils where there was minimal stimulation in CO oxidation rate and carboxydotrophic taxa (Figs. 1, 3). These findings suggest that carboxydotrophic growth of key taxa underlies the enhanced carbon fixation activity in most but not all ecosystem soils.

**Figure 5.**
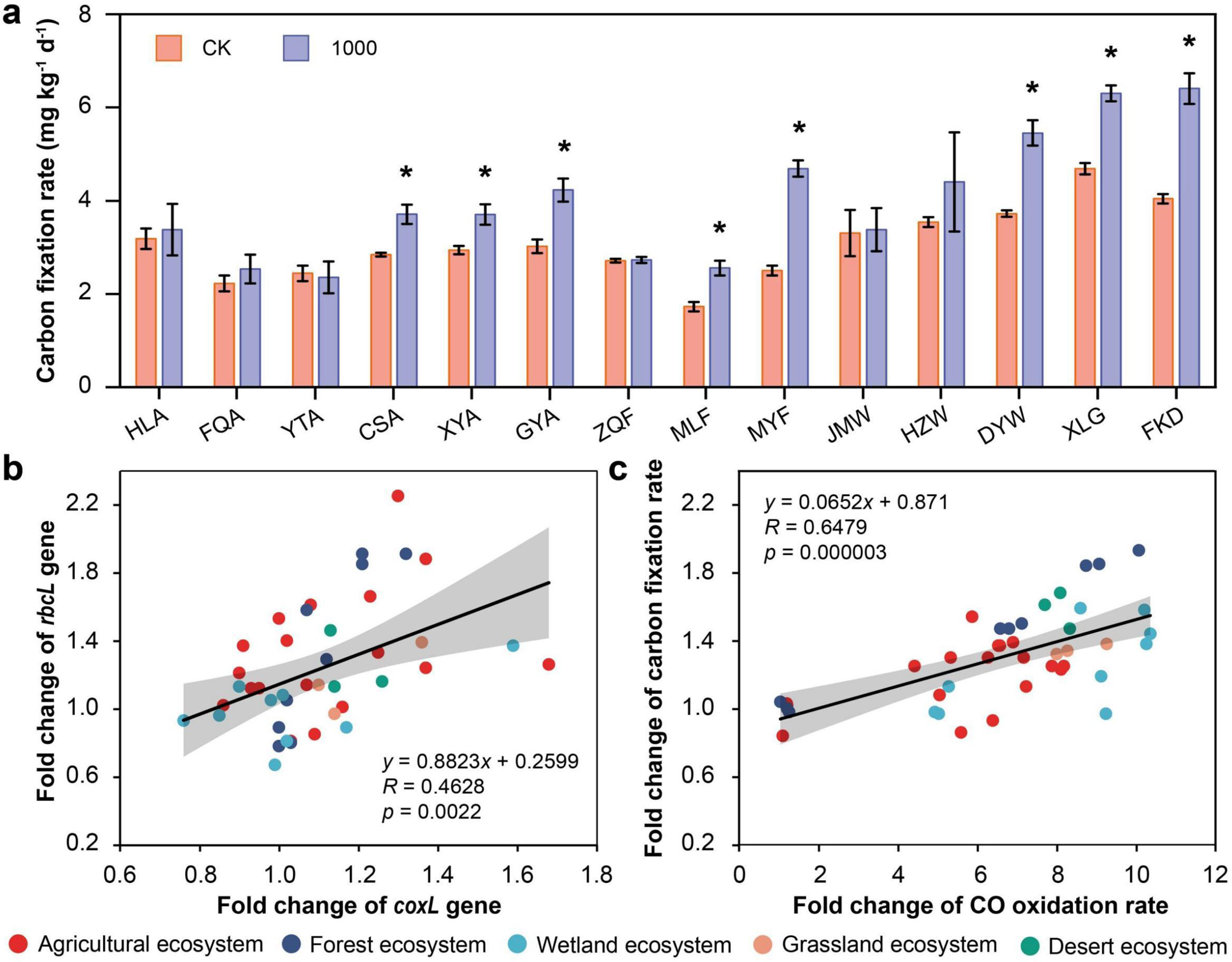
Carbon fixation rate in different treatments after 28 days. **a** Stacked bar chart showing carbon fixation rate in control and 1000 ppmv CO treatment. * indicates significant differences between treatments (two-sided Wilcoxon test). **b** Spearman’s correlations between fold change of *coxL* gene and fold change of *rbcL* gene. **c** Spearman’s correlations between CO oxidation rate and carbon fixation rate. The shaded area in gray represents the 95% confidence interval.

Finally, we explored the correlation of the gene abundance and activity between CO oxidation and carbon fixation in soils after elevated CO treatments. Linear correlation analysis verified that the fold change of genes encoding the CO dehydrogenase (*coxL*) and CO oxidation rate were both significantly increased with the fold change of genes encoding the RuBisCO (*rbcL*) (*r* = 0.46, *p* = 0.0022) and carbon fixation rate (*r* = 0.65, *p* = 0.000003) (Fig. 5b, c). Thus, soil CO oxidizers and oxidation activities likely markedly contribute to soil carbon fixation in the different terrestrial ecosystems. These correlations align with recent studies^54^ indicating that carboxydotrophic bacteria use CO-dehydrogenases as a general mechanism for aerobic respiration and carbon fixation, thereby supporting the productivity of terrestrial ecosystems. These findings strengthen the conclusion that soil carboxydotrophic bacteria is a significant contributor to the global carbon flux^3,7^.

### Conclusions

In summary, our study uncovers a diverse array of CO-oxidizing bacteria, both phylogenetically and physiologically, present in various terrestrial ecosystem soils. While most of these bacteria are organoheterotrophs capable of subsisting on trace amounts of CO, a select few are facultative autotrophs that utilize CO when it is available in higher concentrations. Remarkably, soil CO oxidizers exhibit sensitivity to slight fluctuations in CO concentrations (as low as 1 ppmv), resulting in enhanced CO uptake. This sensitivity underscores the significant role of soils as effective CO sinks. Our microcosm experiments revealed considerable variation in the lineages of bacteria, CO-dehydrogenases, and RuBisCO that became enriched in response to CO availability, reflecting the original community structures of the different soil types. However, certain lineages such as *Bradyrhizobium* and a Pseudonocardiaceae MAG were widespread across diverse soils and strongly stimulated by CO exposure, becoming more abundant and sometimes even the dominant genera in elevated CO treatments. Furthermore, we established a strong correlation between CO oxidation and carbon fixation, indicating that the carboxydotrophic growth of key taxa contributes to enhanced carbon fixation activity in eight of the 14 soils tested. Overall, while CO exposre was previously considered toxic, our findings suggest it does not significantly alter soil properties, microbial abundance, or microbial richness, and instead promotes the activity of CO oxidizers. In turn, the increased prevalence of carboxydotrophic taxa may influence biogeochemical processes, as evidenced by the enhanced genetic potential and biochemical activity for carbon fixation observed in most ecosystem soils. Building on these insights, we anticipate that elevated CO emissions originating from natural or anthropogenic sources would promote the growth of facultative carboxydotrophs, although the specific lineages stimulated are likely to vary across different ecosystem soils. Such lineages would be able to regulate atmospheric CO levels, in turn reducing the toxic and climate-forcing levels of these gases, while potentially contributing to chemosynthetic primary production.

## Methods

### Sample collection, microcosm design and soil sampling

Soil samples for microcosm experiment were collected in 2020 and 2021 from different representative ecosystems at fourteen sites in China based on the Chinese Ecosystem Research Network. Agricultural, forest, wetland, grassland, and desert ecosystems were sampled at 6 (Hailun City in Heilongjiang Province, Fengqiu City in Henan Province, Yingtan City Guiyang City in Guizhou Province), 3 (Zhaoqing City in Guangdong Province, Mengla City in Yunnan Province, and Mianyang City in Sichuan Province), 3 (Jiamusi City in Heilongjiang Province, Hongze City in Jiangsu Province, and Dongying City in Shandong Province), 1 (Xilingol League in Inner Mongolia Autonomous Region), and 1 (Fukang City in Xinjiang Uygur Autonomous Region) sites, respectively (Fig. 1, Details showed in Supplementary Table 1). At each site, topsoil (0-20 cm) was collected aseptically in triplicate from an area of approximately 400 m² and then pooled to create a soil mixture. These soil samples were air-dried in the lab and subsequently sieved through a 10-mesh (2 mm) sieve to eliminate roots and large debris prior to the microcosm experiments. Detailed soil properties were listed in our previous study^17^.

Before setting up the microcosm experiment, the soil moisture content was set to 10% (w/w) and incubated at 30 °C for one week to reactivate the soil microbial community. To investigate the impact of CO supplementation on soil microbial communities and activity, a control (CK: 0 ppmv) and three elevated CO levels (Treatment groups: 1, 100 and 1,000 ppmv) were set up. Briefly, approximately 20 g of soil (dry weight) was transferred to a 120 mL serum bottle, sealed with butyl rubber stoppers, and the water content was adjusted to 60% of the maximum field holding capacity using sterilized water. These bottles were purged with synthetic air (21% O_2_ and 360 ppmv CO_2_ balanced with N_2_, Nanjing Wenda Special Gas Co., Ltd.) for 30 seconds. A specific amount of ultra-pure CO gas (99.995%; Nanjing Wenda Special Gas Co., Ltd.) was introduced to achieve four initial headspace concentrations of CO levels in combination with an appropriate volume of synthetic air. Every day, the control and treatment serum bottles underwent a process of flushing with synthetic air followed by the re-establishment of initial CO concentrations, ensuring a consistent supply of CO. Additionally, to ensure soil moisture remained at 60% of its maximum field holding capacity, water was added every two days as needed. The experimental procedures were performed in triplicate and incubated in the dark at 30 °C for 28 days. Samples were taken on days 0, 7, 14, 21, and 28 for subsequent analysis.

### Soil physicochemical and CO oxidation rate analysis

Air-dried soil samples were passed through a 60-mesh (0.25 mm) sieve to ensure a uniform matrix for the analysis of physicochemical properties. The methods of determination soil pH, soil organic matter content (SOM), total nitrogen (TN), nitrate (NO_3_-N), ammonium (NH_4_-N), available published^17,55^. The specific soil properties in different treatments after 28 days were listed in Supplementary Table 4.

During the microcosm experiment, the CO oxidation activity was measured by a gas chromatographic assay using the previous methods with some modifications^13^. Briefly, a total of 5 g of previously collected soil samples was placed into a 120-mL serum vial, and the static headspace of each serum bottle was first flushed with synthetic air for 3min. Then, a defined volume of CO gas mixture (via 5% v/v CO in N_2_ gas cylinder, 99.995% pure) were injected to obtain an initial concentration of ∼1,000 ppm in the static headspace. After incubating the samples at 30°C in the dark for 4 hours, 2 mL of gas samples were periodically withdrawn from the headspace of the serum vials. These samples were then injected into a gas chromatograph with a reduction gas detector via the injection port for analysis (GC7890A, J&W Scientific; Agilent Technologies). No noteworthy CO uptake was detected in control experiments that utilized sterile soils or empty bottles. The CO oxidation rate was calculated by applying first-order kinetics and expressed as nmol g^-1^ h^-1^ at 1000 ppmv CO.

### DNA extraction and real time PCR analysis

Total community DNA was extracted from 168 soil samples, with 0.5 g of each sample, using the E.Z.N.A.® Stool DNA Kit following the manufacturer’s guidelines (Omega Bio-Tek, Norcross, GA). The extracted DNA’s quality, quantity, and purity were evaluated using electrophoresis on a 0.8% agarose gel and a Nanodrop® ND-1000 UV-Vis spectrophotometer (NanoDrop Technologies, Wilmington, DE). The DNA samples were stored at −80 °C before use.

To estimate the total bacterial biomass of extracted soil samples, we used the quantitative real-time PCR (q-PCR) method with the 515F (5’-GTGCCAGCMGCCGCGG-3’) and 907R (5’-CCGTCAATTCMTTTRAGTTT-3’) primers^41^. q-PCR was performed using a CFX96 Real-Time PCR Detection System (Bio-Rad Laboratories, Inc., Hercules, CA) in a 20 μL reaction mixtures consisted of 1 μL template DNA, 0.4 μL of each primer (10 μM), 10 μL 2× ChamQ Universal SYBR qPCR Master Mix (Vazyme Biotech Co., Ltd., Nanjing, China) and 8.2 μL ddH_2_O. The amplification process involved 40 cycles: denaturation at 95 °C for 30 s, primer annealing at 55 °C for 30 s, and extension at 72 °C for 30 s. Specificity was checked with a melting curve analysis, and negative controls were included to detect contamination. Each sample was analyzed in triplicate, and a standard curve was generated using a 10-fold dilution series of plasmids containing the bacterial 16S rRNA gene from environmental samples. The amplification efficiency ranged from 97.6-99.8%, with R^2^ values of 0.989-0.996.

### 16S rRNA gene sequencing and analysis

The V4-V5 region of the 16S rRNA gene was employed to assess the soil microbial community composition, using the universal prokaryotic primer set 515F/907R across all soil samples, as described in previous studies^56^. Briefly, the DNA samples were amplified and paired-end sequenced by Shanghai Biozeron Biological Technology Co. Ltd. (Shanghai, China) on the Illumina MiSeq PE250 platform (Illumina, San Diego, CA, USA) according to the manufacturer’s protocols. The raw sequences were demultiplexed, quality-filtered using Trimmomatic, and merged with FLASH following established protocols as used in previous studies^41,56^. The merged sequences were subjected to noise reduction with DADA2 in Qiime2, and the taxonomic information of the obtained ASVs (Amplicon Sequence Variant) was identified based on the SILVA database (release 138)^57^. Alpha diversity measures such as observed richness, Chao1 richness, and Shannon indexes were computed using mothur (version 1.30.1) with their default settings^57^.

### Metagenomic sequencing, assembly and binning

Based on the results of taxonomic analysis of 16S rRNA gene amplicon sequencing and CO oxidation rate, samples from higher CO concentration treatment (1,000 ppmv), which showed significant shifts in bacterial communities and CO oxidation activity compared to the control group (CK, 0 ppmv), were selected for metagenomic analysis (84 samples in total). The extracted DNA was fragmented to approximately 450 bp using a Covaris S220 Focused Ultrasonicator (Covaris, Woburn, MA, USA). Metagenomic libraries were prepared and sequenced on the Illumina NovaSeq 6000 platform at Shanghai Biozeron Biological Technology Co. Ltd., generating around 10 Gb of raw data per sample. These raw reads were processed for quality control using fastp (v. 0.23.2, default parameters)^58^ and assembled and binned using the metaWRAP pipeline (v. 1.3.2)^59^. Briefly, the quality-controlled reads were individually assembled and co-assembled using Megahit v. 1.1.3 in the metaWRAP Assembly module (default parameters)^59^. Contigs shorter than 1,000 bp were discarded. Each assembly was binned using the metaWRAP Binning module (parameters: -metabat2 -maxbin2 -concoct). Then, the three bin sets were refined and consolidated with the Bin_refinement module using specific parameters (-c 50, -x 10)^60^. The refined bins were subsequently consolidated and dereplicated to a nonredundant set of species-level metagenome-assembled genomes (MAGs) using dRep v3.4.0, with parameters set to -comp 50 -con 10^61^, ensuring 95% average nucleotide identity (ANI). MAGs were assessed for completeness and contamination with CheckM (v. 1.0.18)^62^, and only those with ≥ 50% completeness and ≤ 10% contamination were retained. In total, we obtained a nonredundant genome set of 630 MAGs (details showed in Supplementary Tables 7, 8). The taxonomic assignments of these MAGs were performed using GTDB-Tk (v. 2.1.1, parameters: -x fa -cpus 66 -full_tree) with the Genome Taxonomy Database (GTDB) (release 05-RS95)^63^. MAG abundance was estimated using CoverM (v. 0.6.0) following the methodology described previously^64^.

### Functional annotation

The metabolic capability of the soil communities was estimated using previously described methods^17,65^. In brief, we constructed a nonredundant gene catalog using CD-HIT (v. 4.8.1, parameters: -c 0.95 -T 0 -M 0 -G 0 -aS 0.9 -g 1 -r 1 -d 0), with representative sequences selected from each cluster for further analysis. Predicted genes from the reference gene catalog were assigned to major metabolic pathways related to aerobic respiration, trace gas metabolism, carbon fixation, sulfur metabolism, nitrogen metabolism and alternative electron donor/acceptor processes using DIAMOND (v. 2.0.8.146, parameters: query-outfmt, 6; threads, 8; max-target-seqs, 1) against reference datasets as detailed previously described^32^. Contigs in MAGs were also annotated using above methods. The identity threshold of key metabolic markers for metagenomic contigs and MAGs has been detailed showed in previous study^32^. To estimate the proportion of community members encoding each metabolic gene, we calculated the transcripts per million (TPM) for each gene and normalized it by dividing by the average TPM across 14 universal single-copy ribosomal marker genes (Supplementary Table 16). Similar function genes were grouped together, and their values were summed to 100%, as previously described^17,65^.

### Phylogenetic analysis

To investigate the distribution and diversity of CO oxidizers and carbon-fixing microbes, phylogenetic trees were constructed for the large subunits of form I CO-dehydrogenase (CoxL) and RuBisCO (RbcL). The details of the phylogenetic tree construction of MAGs were presented in our previous study^41^. Protein sequences were retrieved from the MAGs and metagenomic contigs via homology-based searches against a subset of reference sequences from a custom database^32^. These sequences were aligned using MUSCLE (v. 5.1)^66^ and further refined with TrimAl (v. 1.2.59)^67^. Maximum-likelihood trees were built using IQ-TREE (v1.6.12)^68^ with best-fit models, bootstrapped with 1000 replicates, and rooted at mid-point. The resulting file was then uploaded to iTOL^69^ for visualization and annotation.

### 13CO2 fixation assays

To test the effect of elevated CO on the capacity of soil carbon fixation, ^13^CO_2_ fixation assays was conducted at the end of the experiment according to a previous study^70^. Briefly, 10 g fresh soil was introduced into a 120-mL serum vial, which was then sealed using a butyl rubber plug and an aluminum cap. The vial was flushed with synthetic air for 3 min, followed by the introduction of either 5% (v/v) ^13^CO2 (99 atom %, Cambridge Isotope Laboratories, Inc., Tewksbury, MA) or 5% (v/v) ^12^CO2, depending on the experimental or control group. The vial was placed in a dark environment at 30 °C for 9 days. The soils were replenished with water every day using the weighing method, and the 5% (v/v) ^13^CO_2_ or ^12^CO_2_ was subsequently reinjected into the vial every two days. Following the incubation period, the soil samples were promptly freeze-dried and treated with 2 M HCl for 24 h. The δ^13^C atom% value of the soil in the vial was determined using a stable isotope ratio mass spectrometer (Delta V Plus, Thermo Fisher Scientific, USA). The carbon fixation rate was calculated based on the differences in ^13^C atom % in the soil before and after incubation and expressed as mg kg^-1^ dry soil d^-1^, following the previous method described by Zhao et al^70^.

### Statistical analysis

All data are expressed as the average ± standard deviation from three independent replicates. Either an unpaired Student’s t-test or two-sided Wilcoxon test was used to compare the means of different treatments. A one-way ANOVA was conducted, followed by Duncan’s multiple range test (*p* < 0.05), to examine significant variations in CO oxidation rate and community structure across various CO concentration treatments using the SPSS 16.0 software package (SPSS, Chicago, IL). Results with *p*-values of \**p* < 0.05 and \*\**p* < 0.01 are considered statistically significant.

## Data availability

All data necessary for our analysis are freely accessible, and their sources are provided in the “Methods” section and the Supplementary Information. The 16S rRNA gene amplicon sequences and the metagenomes have been deposited in the Sequence Read Archive (SRA) of the NCBI with accession numbers PRJNA1173976, PRJNA1174050 and PRJNA1175901, respectively.

## Acknowledgements

This research was supported by the National Natural Science Foundation of China (42130718, 42207030, 41991335), the National Key Research and Development Program of China (2019YFC1803705), the Jiangsu Funding Program for Excellent Postdoctoral Talent (2022ZB465), the China Postdoctoral Science Foundation (2022M723241), and the National Science and Technology Innovation Leading Talents Program (SQ2022RA24910167). P.M.L acknowledges an ARC DECRA Fellowship (DE250101210) for salary support. C.G. acknowledges an ARC DECRA Fellowship for project development (DE170100310) and NHMRC EL2 Fellowship (APP1178715) for salary support.

## Author contributions

C.G. Y.X. and Y.T. conceived this study. Y.T. and C.G. supervised this study. Y.X. and Y.T. performed experiments and data analysis. Y.T. J.L. P.M.L. X.D. and C.G. assisted with meta-omic analysis. X.P. and Z.G. assisted with field soil sample collection. S.D. Y.S. W.H. H.W. Y.L. Y.H. and Y.L. provided critical comments on this study. Y.X. Y.T. and C.G. wrote the paper with input from all authors.

## References

1 Oelgeschläger, E. & Rother, M. Carbon monoxide-dependent energy metabolism in anaerobic bacteria and archaea. Arch. Microbiol. 190, 257–269 (2008).

2 Sobieraj, K., Stegenta-Dabrowska, S., Luo, G., Koziel, J. A. & Bialowiec, A. Carbon monoxide fate in the environment as an inspiration for biorefinery industry: A review. Front. Env. Sci. 10, 822463 (2022).

3 Greening, C. & Grinter, R. Microbial oxidation of atmospheric trace gases. Nat. Rev. Microbiol. 20, 513–528 (2022).

4 Kleiner, M. et al. Metaproteomics of a gutless marine worm and its symbiotic microbial community reveal unusual pathways for carbon and energy use. P. Natl. Acad. Sci. USA 109, E1173–E1182 (2012).

5 Alonso, J. R., Cardellach, F., López, S., Casademont, J. & Miró, O. Carbon monoxide specifically inhibits cytochrome C oxidase of human mitochondrial respiratory chain. Pharmacol. Toxico. 93, 142–146 (2003).

6 Hopper, C. P. et al. Role of Carbon Monoxide in Host-Gut Microbiome Communication. Chem. Rev. 120, 13273–13311 (2020).

7 King, G. M. & Weber, C. F. Distribution, diversity and ecology of aerobic CO-oxidizing bacteria. Nat. Rev. Microbiol. 5, 107–118 (2007).

8 Miyakawa, S., Yamanashi, H., Kobayashi, K., Cleaves, H. J. & Miller, S. L. Prebiotic synthesis from CO atmospheres: implications for the origins of life. P. Natl. Acad.Sci. 99, 14628–14631 (2002).

9 Wang, J. J. et al. CO-driven electron and carbon flux fuels synergistic microbial reductive dechlorination. Microbiome 12, 154 (2024).

10 Tiquia-Arashiro, S. M. & Tiquia-Arashiro, S. M. CO-oxidizing Microorganisms. Thermophilic carboxydotrophs and their applications in biotechnology, 11–28 (2014).

11 Ji, M. et al. Atmospheric trace gases support primary production in Antarctic desert surface soil. Nature 552, 400–403 (2017).

12 Conrad, R. Soil microorganisms as controllers of atmospheric trace gases (H_2_, CO, CH_4_, OCS, N_2_O, and NO). Microbiol. Rev. 60, 609–640 (1996).

13 Cordero, P. R. et al. Atmospheric carbon monoxide oxidation is a widespread mechanism supporting microbial survival. ISME J. 13, 2868–2881 (2019).

14 Bayly, K. et al. Mycobacteria tolerate carbon monoxide by remodeling their respiratory chain. Msystems 6, e01292–20 (2021).

15 Bay, S. K. et al. Trace gas oxidizers are widespread and active members of soil microbial communities. Nat. Microbiol. 6, 246–256 (2021).

16 de la Porte, A., Durand, A.-A., Whalen, J., Yergeau, É. & Constant, P. A rhizosphere effect promotes the persistence of gas oxidization activity in soil. Soil Biol. Biochem. 199, 109599 (2024).

17 Xu, Y. et al. Atmospheric trace gas oxidizers contribute to soil carbon fixation driven by key soil conditions in terrestrial ecosystems. Environ. Sci. Technol. 58, 21617–21628 (2024).

18 King, G. M. Molecular and culture-based analyses of aerobic carbon monoxide oxidizer diversity. Appl. Environ. Microbiol. 69, 7257–7265 (2003).

19 Cunliffe, M. Correlating carbon monoxide oxidation with cox genes in the abundant marine Roseobacter clade. ISME J. 5, 685–691 (2011).

20 King, G. M. Uptake of carbon monoxide and hydrogen at environmentally relevant concentrations by mycobacteria. Appl. Environ. Microbiol. 69, 7266–7272 (2003).

21 Islam, Z. F. et al. Two Chloroflexi classes independently evolved the ability to persist on atmospheric hydrogen and carbon monoxide. ISME J. 13, 1801–1813 (2019).

22 King, C. E. Diversity and activity of aerobic thermophilic carbon monoxide-oxidizing bacteria on Kilauea Volcano, Hawaii. (Louisiana State University and Agricultural & Mechanical College, 2013).

23 McDuff, S., King, G., Neupane, S. & Myers, M. Isolation and characterization of extremely halophilic CO-oxidizing Euryarchaeota from hypersaline cinders, sediments and soils and description of a novel CO oxidizer, *Haloferax namakaokahaiae* Mke2. 3T, sp. nov. FEMS Microbiol. Ecol. 92, fiw028 (2016).

24 Weber, C. F. & King, G. M. The phylogenetic distribution and ecological role of carbon monoxide oxidation in the genus Burkholderia. FEMS Microbiol. Ecol. 79, 167–175 (2012).

25 Weber, C. F. & King, G. M. Volcanic soils as sources of novel CO-oxidizing *Paraburkholderia* and *Burkholderia*: *Paraburkholderia hiiakae* sp. nov., *Paraburkholderia metrosideri* sp. nov., *Paraburkholderia paradisi* sp. nov., *Paraburkholderia peleae* sp. nov., Microbiol. 8, 207 (2017).

26 Sokolova, T. et al. Aerobic carbon monoxide oxidation in the course of growth of a hyperthermophilic archaeon, Sulfolobus sp. ETSY. Microbiology 86, 539–548 (2017).

27 Krüger, B. & Meyer, O. Thermophilic Bacilli growing with carbon monoxide. Arch. Microbiol. 139, 402–408 (1984).

28 Lorite, M. a. J., Tachil, J. r., Sanjuan, J. n., Meyer, O. & Bedmar, E. J. Carbon monoxide dehydrogenase activity in Bradyrhizobium japonicum. Appl. Environ. Microbiol. 66, 1871–1876 (2000).

29 Novelli, P., Masarie, K. & Lang, P. Distributions and recent changes of carbon monoxide in the lower troposphere. J. Geophys. Res. Atmos. 103, 19015–19033 (1998).

30 Khalil, M. & Rasmussen, R. The global cycle of carbon monoxide: Trends and mass balance. Chemosphere 20, 227–242 (1990).

31 Bartholomew, G. W. & Alexander, M. Soil as a sink for atmospheric carbon monoxide. Science 212, 1389–1391 (1981).

32. Ortiz, M., et al. Multiple energy sources and metabolic strategies sustain microbial diversity in Antarctic desert soils. P. Natl. Acad. Sci. USA 118, e2025322118 (2021).

33 Granier, C. et al. Evolution of anthropogenic and biomass burning emissions of air pollutants at global and regional scales during the 1980–2010 period. Climatic Change 109, 163–190 (2011).

34 King, G. M. & Crosby, H. Impacts of plant roots on soil CO cycling and soil–atmosphere CO exchange. Global Change Biol. 8, 1085–1093 (2002).

35 Barnard, R. J. & Weber, J. S. Carbon monoxide: a hazard to fire fighters. Arch. Environ.l Health 34, 255–257 (1979).

36 Ou-Yang, C.-F. et al. Characteristics of atmospheric carbon monoxide at a high-mountain background station in East Asia. Atmos. Environ. 89, 613–622 (2014).

37 Palmer, J. L., Hilton, S., Picot, E., Bending, G. D. & Schäfer, H. Tree phyllospheres are a habitat for diverse populations of CO-oxidizing bacteria. Environ. Microbiol. 23, 6309–6327 (2021).

38 Denton, M. D., Reeve, W. G., Howieson, J. G. & Coventry, D. R. Competitive abilities of common field isolates and a commercial strain of *Rhizobium leguminosarum* bv. trifolii for clover nodule occupancy. Soil Biol. Biochem. 35, 1039–1048 (2003).

39 Laguerre, G., Louvrier, P., Allard, M.-R. & Amarger, N. Compatibility of rhizobial genotypes within natural populations of Rhizobium leguminosarum biovar viciae for nodulation of host legumes. Appl. Environ. Microbiol. 69, 2276–2283 (2003).

40 Hendrickson, O. & Kubiseski, T. Soil microbial activity at high levels of carbon monoxide. Report No. 0047-2425, (Wiley Online Library, 1991).

41 Xu, Y. et al. Genome-resolved metagenomics reveals how soil bacterial communities respond to elevated H2 availability. Soil Biol. Biochem. 163, 108464 (2021).

42 Fuglestvedt, J., Isaksen, I. & Wang, W.-C. Estimates of indirect global warming potentials for CH_4_, CO and NO_x_. Climatic Change 34, 405–437 (1996).

43 Delgado-Baquerizo, M. et al. A global atlas of the dominant bacteria found in soil. Science 359, 320–325 (2018).

44 Janssen, P. H. Identifying the dominant soil bacterial taxa in libraries of 16S rRNA and 16S rRNA genes. Appl. Environ. Microbiol. 72, 1719–1728 (2006).

45 Brown, C. T. et al. Unusual biology across a group comprising more than 15% of domain Bacteria. Nature 523, 208–211 (2015).

46 Giguere, A. T. et al. Acidobacteria are active and abundant members of diverse atmospheric H2-oxidizing communities detected in temperate soils. ISME J. 15, 363–376 (2021).

47 Hardy, K. R. & King, G. M. Enrichment of high-affinity CO oxidizers in Maine forest soil. Appl. Environ. Microbiol. 67, 3671–3676 (2001).

48 Conrad, R. & Seiler, W. Utilization of traces of carbon monoxide by aerobic oligotrophic microorganisms in ocean, lake and soil. Arch. Microbiol. 132, 41–46 (1982).

49 Conrad, R., Meyer, O. & Seiler, W. Role of carboxydobacteria in consumption of atmospheric carbon monoxide by soil. Appl. Environ. Microbiol. 42, 211–215 (1981).

50 Park, S. W., Park, S. T., Lee, J. E. & Kim, Y. M. Pseudonocardia carboxydivorans sp. nov., a carbon monoxide-oxidizing actinomycete, and an emended description of the genus Pseudonocardia. Int. J. Syst. Evol. Micr. 58, 2475–2478 (2008).

51 King, C. & King, G. Description of Thermogemmatispora carboxidivorans sp. nov., a carbon-monoxide-oxidizing member of the class Ktedonobacteria isolated from a geothermally heated biofilm, and analysis of carbon monoxide oxidation by members of the class Ktedonobacteria. Int. J. Syst. Evol. Micr. 64, 1244–1251 (2014).

52 Jaffe, A. L. et al. Saccharibacteria harness light energy using type-1 rhodopsins that may rely on retinal sourced from microbial hosts. ISME J. 16, 2056–2059 (2022).

53 Cross, K. L. et al. Targeted isolation and cultivation of uncultivated bacteria by reverse genomics. Nat. Biotechnol. 37, 1314–1321 (2019).

54 Lappan, R. et al. Molecular hydrogen in seawater supports growth of diverse marine bacteria. Nat. Microbiol. 8, 581–595 (2023).

55 Xu, Y. et al. Occurrence and risk assessment of potentially toxic elements and typical organic pollutants in contaminated rural soils. Sci. Total Environ. 630, 618–629 (2018).

56 Xu, Y. et al. Endogenous biohydrogen from a rhizobium-legume association drives microbial biodegradation of polychlorinated biphenyl in contaminated soil. Environ. Int. 176, 107962 (2023).

57 Bolyen, E. et al. Reproducible, interactive, scalable and extensible microbiome data science using QIIME 2. Nat. Biotechnol. 37, 852–857 (2019).

58 Uritskiy, G. V., DiRuggiero, J. & Taylor, J. MetaWRAP-a flexible pipeline for genome-resolved metagenomic data analysis. Microbiome 6, 1–13 (2018).

59 Li, D., Liu, C.-M., Luo, R., Sadakane, K. & Lam, T.-W. MEGAHIT: an ultra-fast single-node solution for large and complex metagenomics assembly via succinct de Bruijn graph. Bioinformatics 31, 1674–1676 (2015).

60 Dong, X. et al. Phylogenetically and catabolically diverse diazotrophs reside in deep-sea cold seep sediments. Nat. Commun. 13, 4885 (2022).

61 Olm, M. R., Brown, C. T., Brooks, B. & Banfield, J. F. dRep: a tool for fast and accurate genomic comparisons that enables improved genome recovery from metagenomes through de-replication. ISME J. 11, 2864–2868 (2017).

62 Parks, D. H., Imelfort, M., Skennerton, C. T., Hugenholtz, P. & Tyson, G. W. CheckM: assessing the quality of microbial genomes recovered from isolates, single cells, and metagenomes. Genome Res. 25, 1043–1055 (2015).

63 Dong, X. et al. Functional diversity of microbial communities in inactive seafloor sulfide deposits. FEMS Microbiol. Ecol. 97, fiab108 (2021).

64 Dong, X. et al. Thermogenic hydrocarbon biodegradation by diverse depth-stratified microbial populations at a Scotian Basin cold seep. Nat. Commun. 11, 5825 (2020).

65 Dong, X. et al. Metagenomic views of microbial communities in sand sediments associated with coral reefs. Microb. Ecol. 85, 465–477 (2023).

66 Edgar, R. C. MUSCLE: a multiple sequence alignment method with reduced time and space complexity. BMC Bioinformatics 5, 1–19 (2004).

67 Capella-Gutiérrez, S., Silla-Martínez, J. M. & Gabaldón, T. trimAl: a tool for automated alignment trimming in large-scale phylogenetic analyses. Bioinformatics 25, 1972–1973 (2009).

68 Nguyen, L.-T., Schmidt, H. A., Von Haeseler, A. & Minh, B. Q. IQ-TREE: a fast and effective stochastic algorithm for estimating maximum-likelihood phylogenies. Mol. Biol. Evol. 32, 268–274 (2015).

69 Letunic, I. & Bork, P. Interactive tree of life (iTOL) v3: an online tool for the display and annotation of phylogenetic and other trees. Nucleic Acids Res. 44, W242–W245 (2016).

70 Zhao, K. et al. Desert and steppe soils exhibit lower autotrophic microbial abundance but higher atmospheric CO_2_ fixation capacity than meadow soils. Soil Biol. Biochem. 127, 230–238 (2018).

