## Supporting Information for "Carbon monoxide exposure stimulates growth and activity of primary producers in diverse soil ecosystems"

### Supplementary Information: list of material provided

#### Supplementary Figures

**Fig. S1.** The relationship between CO mixing ratio, soil physicochemical properties, and beta diversity (genus level) are visualised by db-RDA.

**Fig. S2.** Maximum-likelihood phylogenetic tree illustrating the key enzymes involved in carbon fixation via the CBB cycle.

**Fig. S3.** Changes of metabolic potential of the microbial communities in representative terrestrial ecosystem soils.

**Fig. S4.** Distribution of metabolic genes of the microbial communities in representative terrestrial ecosystem soils.

#### Supplementary Tables

**Table S1.** Description of soil sampling sites in different ecosystems used in this study.

**Table S2.** Abundance, diversity and composition of the microbial communities in different treated soil samples.

**Table S3.** Soil physicochemical properties in different treatments after 28 days.

**Table S4.** Results of marginal permutation tests of db-RDA for Fig. S1.

**Table S5.** Relative abundance of the taxa at the genus level based on 16S rRNA gene amplicon sequencing at different treated soil samples.

**Table S6.** Relative abundance of the taxa at the phylum level based on 16S rRNA gene amplicon sequencing at different treated soil samples.

**Table S7.** Summary statistics of archaeal and bacterial MAGs assembled from soil samples.

**Table S8.** METABOIC annotation results for 630 MAGs.

**Table S9.** Relative abundance of each MAG (%).

**Table S10.** Dynamic changes of CO oxidation rate (nmol $g^{-1}$ $h^{-1}$) in different treatments.

**Table S11.** Metabolic potential of the microbial communities in soil samples.

**Table S12.** Summary of metagenome-assembled genomes.

**Table S13.** Number of key metabolic marker genes in MAG.

**Table S14.** Presence / Absence of key metabolic marker genes in MAG.

**Table S15.** Carbon fixation rate (mg kg(dry weight)$^{-1}$ d$^{-1}$) at elevated CO treatments in different ecosystem soils.

**Table S16.** Abundance of 14 marker genes in soil samples. Gene are represented in units of transcripts per million (TPM).

### Supplementary Figures

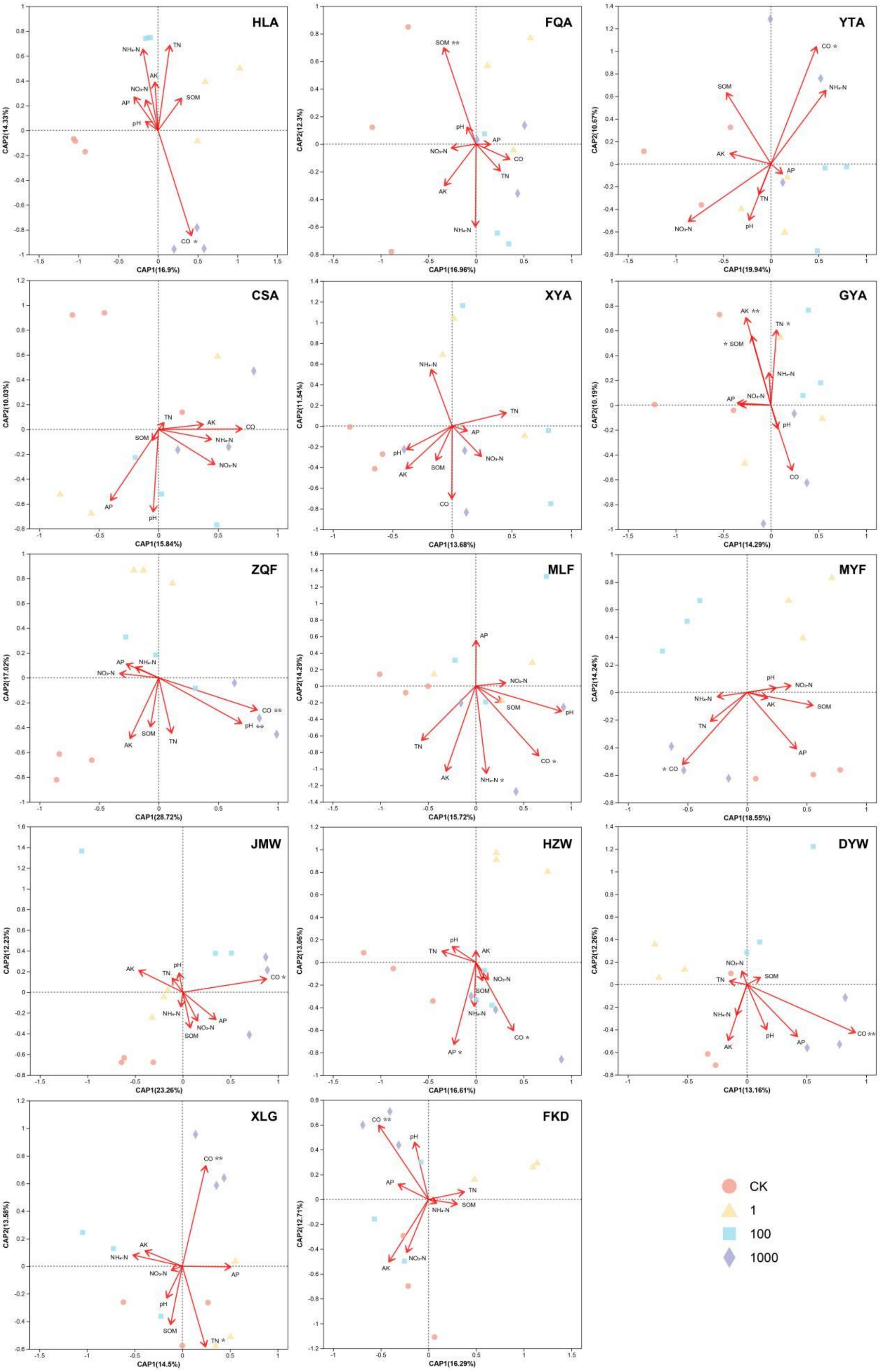

**Fig. S1. The relationship between CO mixing ratio, soil physicochemical properties, and beta diversity (genus level) are visualised by db-RDA.** *p* values are denoted by asterisks (* $p < 0.05$, ** $p < 0.01$). Results of marginal permutation tests of db-RDA are shown in Table S5. The abbreviations SOM, TN, $NO_3$-N, $NH_4$-N, AP, and AK referred to soil organic matter, total nitrogen, nitrate nitrogen, ammonium nitrogen, available phosphorus, and available potassium, respectively.

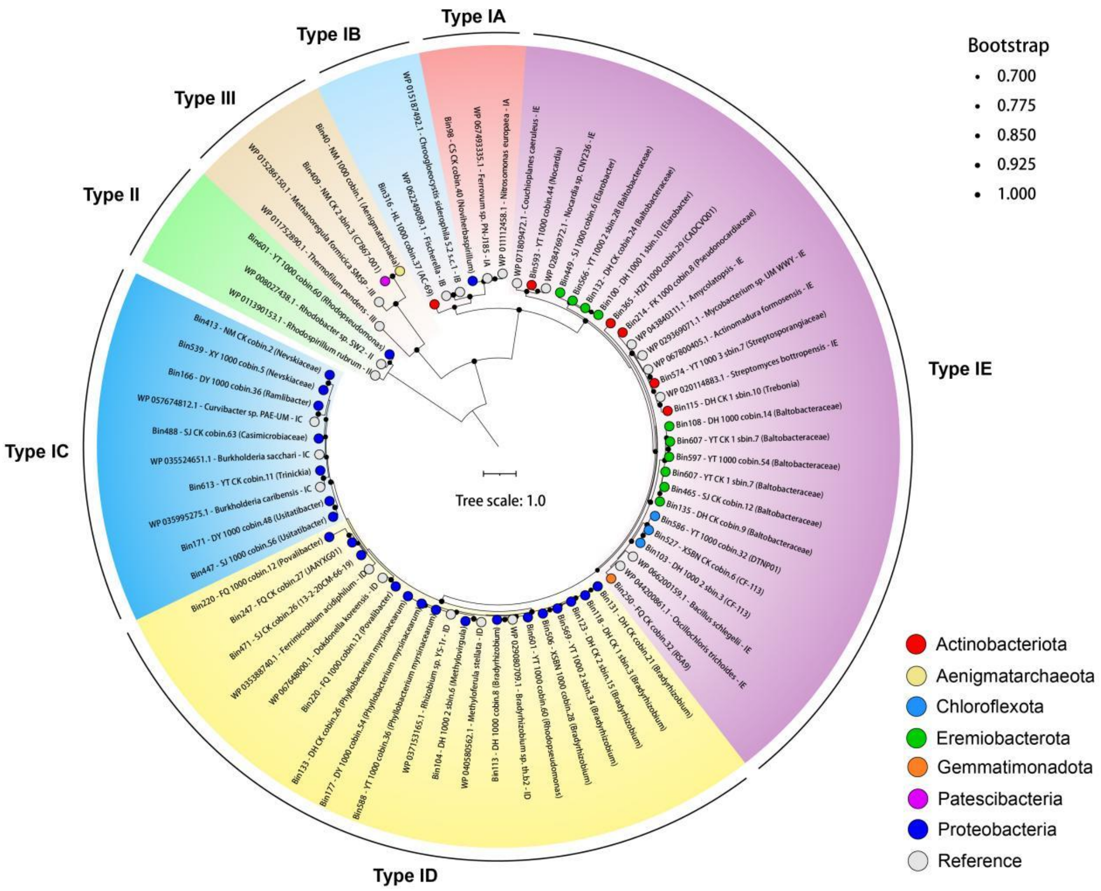


**Fig. S2. Maximum-likelihood phylogenetic tree illustrating the key enzymes involved in carbon fixation via the CBB cycle.** The tree displays the amino acid sequences of the large subunits of RuBisCO. Tree was rooted at mid-point while node support (1000 ultrafast bootstrap replicates) was shown in black dots. The colored leaf nodes denote phylum-level taxonomy of MAGs sequences.

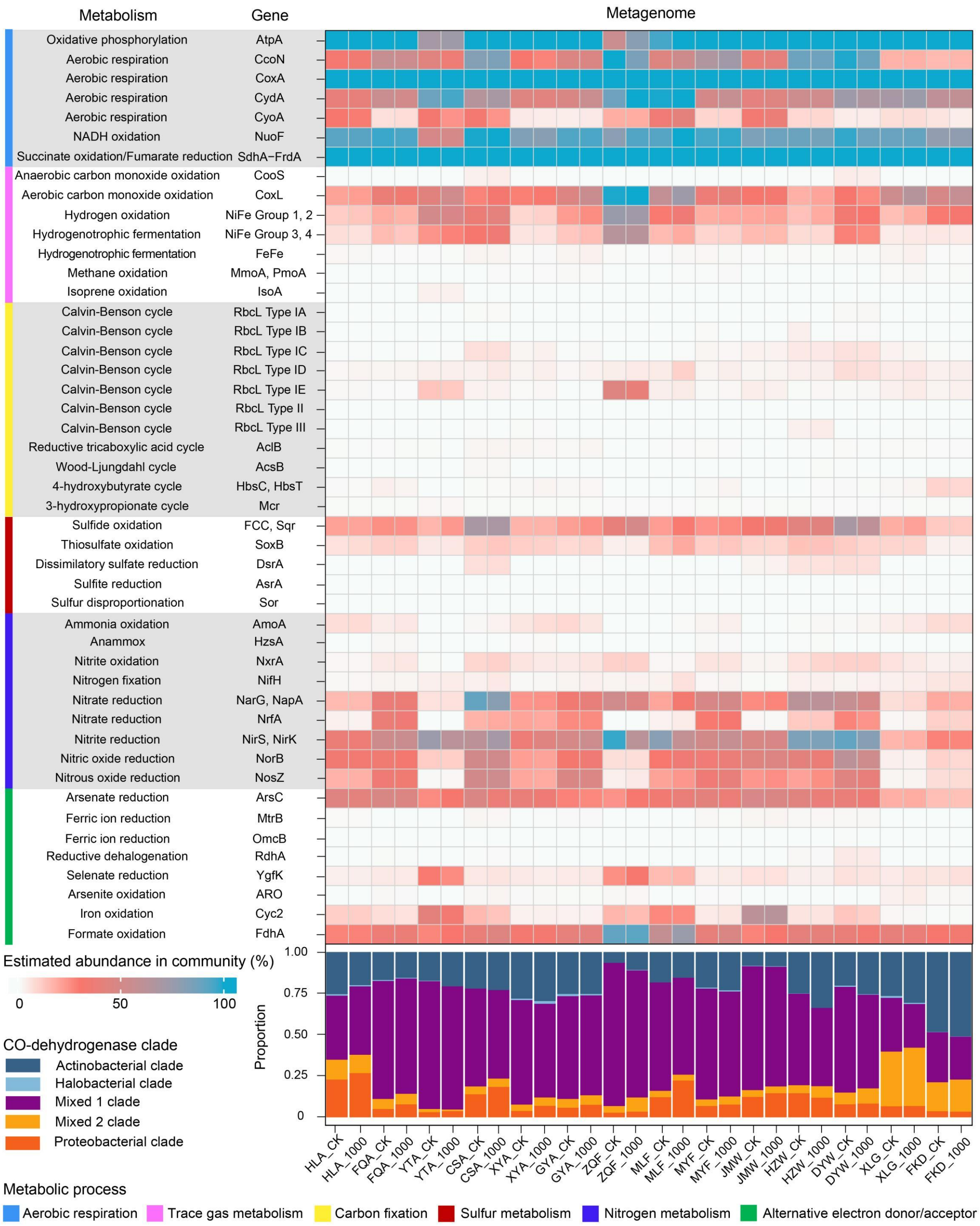


**Fig. S3. Changes of metabolic potential of the microbial communities in representative terrestrial ecosystem soils.** To infer gene abundance in metagenomes, read counts were normalized to gene length and the abundance of 14 single-copy marker genes. Stacked bar chart showing the proportion of different clade in CO-dehydrogenase.

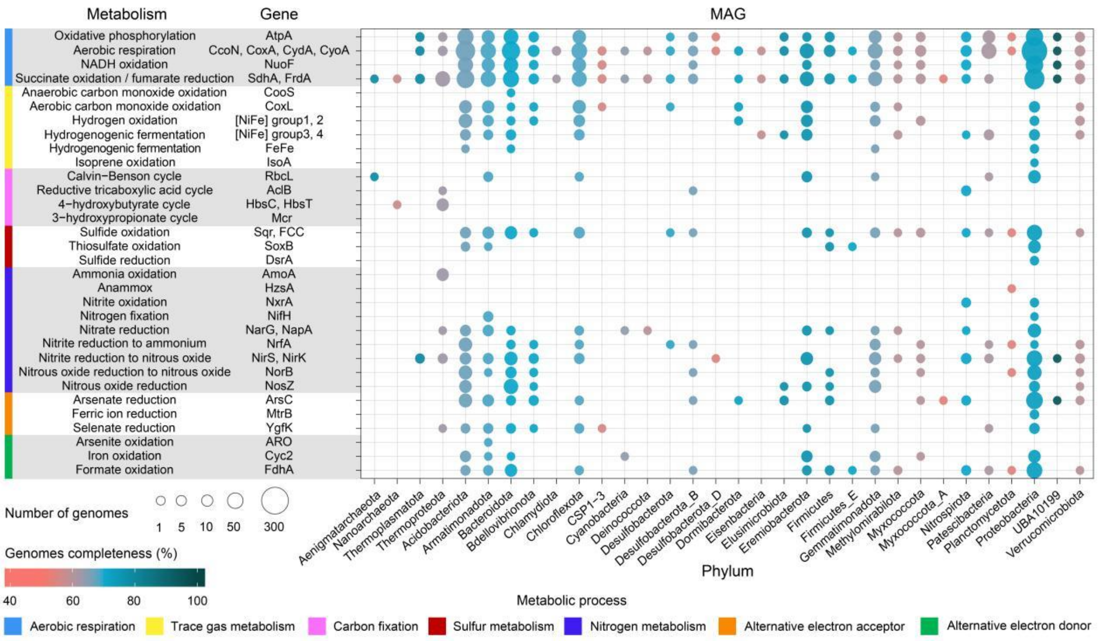


**Fig. S4. Distribution of metabolic genes of the microbial communities in representative terrestrial ecosystem soils.** Dot plot showing the metabolic potential of the 630 MAGs. The magnitude of each data point is indicative of the quantity of genomes within each taxonomic group that contain the specified gene, while the color intensity signifies the mean level of genome completeness.
